# Mass spectrometric analysis of free methionine oxidation levels in *E. coli*

**DOI:** 10.64898/2026.05.12.724599

**Authors:** Maria Ahmed, Philip Bellomio, Bruno Manta, Kyle Swovick, Kevin A. Welle, Jennifer R. Hryhorenko, Sina Ghaemmaghami

## Abstract

Oxidation of free methionine plays important roles in cellular redox homeostasis, yet its accurate quantification has been hindered by methodological challenges. Here, we introduce free Methionine Oxidation by Blocking (fMObB), a mass spectrometry–based method that enables accurate measurement of the fractional oxidation of free methionines. Applying fMObB to *Escherichia coli*, we quantify free methionine oxidation under basal and oxidative stress conditions, and in strains lacking methionine sulfoxide reductases. We find that during oxidative stress, free methionines exhibit higher oxidation levels than protein-bound methionines and that methionine sulfoxide reductases play a central role in maintaining reduced free methionine pools. Together, this work establishes fMObB as a generalizable strategy for probing free methionine redox states in cellular systems.

## Introduction

Within cells, methionines (Mets) can be oxidized to methionine sulfoxides (MetOs) by reactions with reactive oxygen species (ROS) or by enzymatic oxidation catalyzed by monooxygenases.^1^ MetOs can subsequently be converted back to Mets by a conserved class of reducing enzymes known as methionine sulfoxide reductases (MSRs).^2,3^ Oxidation of protein-bound Met residues frequently leads to protein misfolding and has been associated with several neurodegenerative diseases and pathogenic aging.^4–6^ Beyond their role in protein damage and repair, cycles of oxidation and reduction of Met residues contribute broadly to cellular redox homeostasis by functioning as a ROS scavenging mechanism. In this context, Met oxidation followed by MSR-mediated reduction consumes NADPH to eliminate ROS, allowing Mets to serve as a sink for reactive oxidants that would otherwise be damaging to other essential cellular components.^7,8^ Moreover, Met oxidation has been shown to act as a regulatory switch for certain oxidative-responsive transcription factors and modulate the activity of a number of enzymes.^9,10^

In comparison to protein-bound Met residues, the oxidation and reduction of free Mets have received less attention. This is despite the fact that free Mets can be as oxidizable as their protein-bound counterparts and that such oxidation can play a critical role in cellular signaling and disease.^11,12,13^ Although most MSRs, including MSRA and MSRB, preferentially reduce protein-bound MetOs, they can also act on free MetOs with reduced efficiencies.^14,15^ Additionally, some bacteria encode specialized MSRs dedicated to reducing free MetO.^16,17^ Free MetOs are biologically important metabolites that can function as scavengers similar to their protein-bound counterparts, serve as electron acceptors for the respiratory chain under anaerobic conditions, act as signaling molecules, and provide precursors for methionine biosynthesis.^18,19^

Despite these important roles, with a few exceptions^20–22^, the oxidation levels of free Met pools have not been systematically quantified *in vivo*. Although Mets and MetOs are readily detectable by mass spectrometry, accurate quantification of their relative *in vivo* abundances presents several technical challenges that have limited their analysis. First, Mets are highly susceptible to oxidation during sample handling and analysis, making it difficult to distinguish MetOs formed *in vivo* from those generated artifactually during mass spectrometric workflows.^23–25^ Second, differences in ionization efficiencies between Mets and MetOs preclude direct measurement of fractional oxidation from their respective signal intensities alone, necessitating the use of spike-in standards or alternative normalization strategies.^20,26,27^

To address these limitations, a method termed Methionine Oxidation by Blocking (MobB) has recently been developed to facilitate the mass spectrometric analysis of Met oxidation levels.^6,25,28^ In this approach, unoxidized Mets are fully oxidized *in vitro* using ^18^O-labeled hydrogen peroxide, and subsequent mass spectrometric measurements of ^18^O- and ^16^O-labeled methionine sulfoxides are used to determine *in vivo* fractional oxidation levels.^25^ Although MobB has been successfully applied to quantify Met oxidation within individual proteins and in large-scale proteomic studies^6,25^, it has not been previously used to measure oxidation levels of free Mets.

Here, we describe a modified version of the MobB methodology, termed free Methionine Oxidation by Blocking (fMObB), that enables quantitative measurement of the fractional oxidation of free Mets. As a first application of this approach, we quantified free Met oxidation levels in the cytosol of *E. coli* under basal and oxidative stress conditions. The results provide insights into methionine redox homeostasis in *E. coli* and establish fMObB as a generalizable strategy for assessing free methionine oxidation levels in diverse cellular systems.

## Results and Discussion

### fMObB can accurately quantify fractional oxidation of free Mets

The detailed experimental workflow of fMObB is provided in the Methods section. Briefly, cellular extracts containing free Mets and MetOs are treated with ^18^O-labeled hydrogen peroxide, stoichiometrically converting all unoxidized Mets to ^18^O-labeled MetOs. Endogenously formed ^16^O-MetOs and *in vitro*-generated ^18^O-MetOs differ in mass but exhibit identical chromatographic retention times and ionization efficiencies. Consequently, relative intensities of their mass spectrometric signals provide quantitative measures of fractional Met oxidation within the extract.

To assess the quantitative accuracy of fMObB, we first generated defined mixtures of free ^16^O- and ^18^O-MetOs and experimentally measured their relative abundances (**Fig. 1A**). ^16^O-MetO was obtained commercially, while ^18^O-MetO was prepared in-house by complete oxidation of Mets using ^18^O-labeled hydrogen peroxide. The oxidation and isotopic purity of both stock solutions was confirmed by liquid chromatography-mass spectrometry (LC-MS) (**Fig. 1B**). The two stock solutions were mixed at defined ratios and relative abundances of ^16^O- and ^18^O-MetOs were quantified in triplicate samples (**Fig. 1C**). The experimentally measured ratios showed excellent agreement with expected values (slope = 0.97, R^2^ >0.99), validating the overall quantitative accuracy of fMObB measurements. However, our experiments detected a background level of ~2% ^16^O-MetO even in samples expected to contain only ^18^O-MetO. This background likely arises from residual oxidation of the commercially obtained Met used to synthesize ^18^O-MetO, or from isotopic impurity in the ^18^O-labeled hydrogen. Depending on which source predominates, the fractional oxidation values reported in the following studies may need to be corrected by 0–2% to reflect true *in vivo* levels.

**Figure 1.**
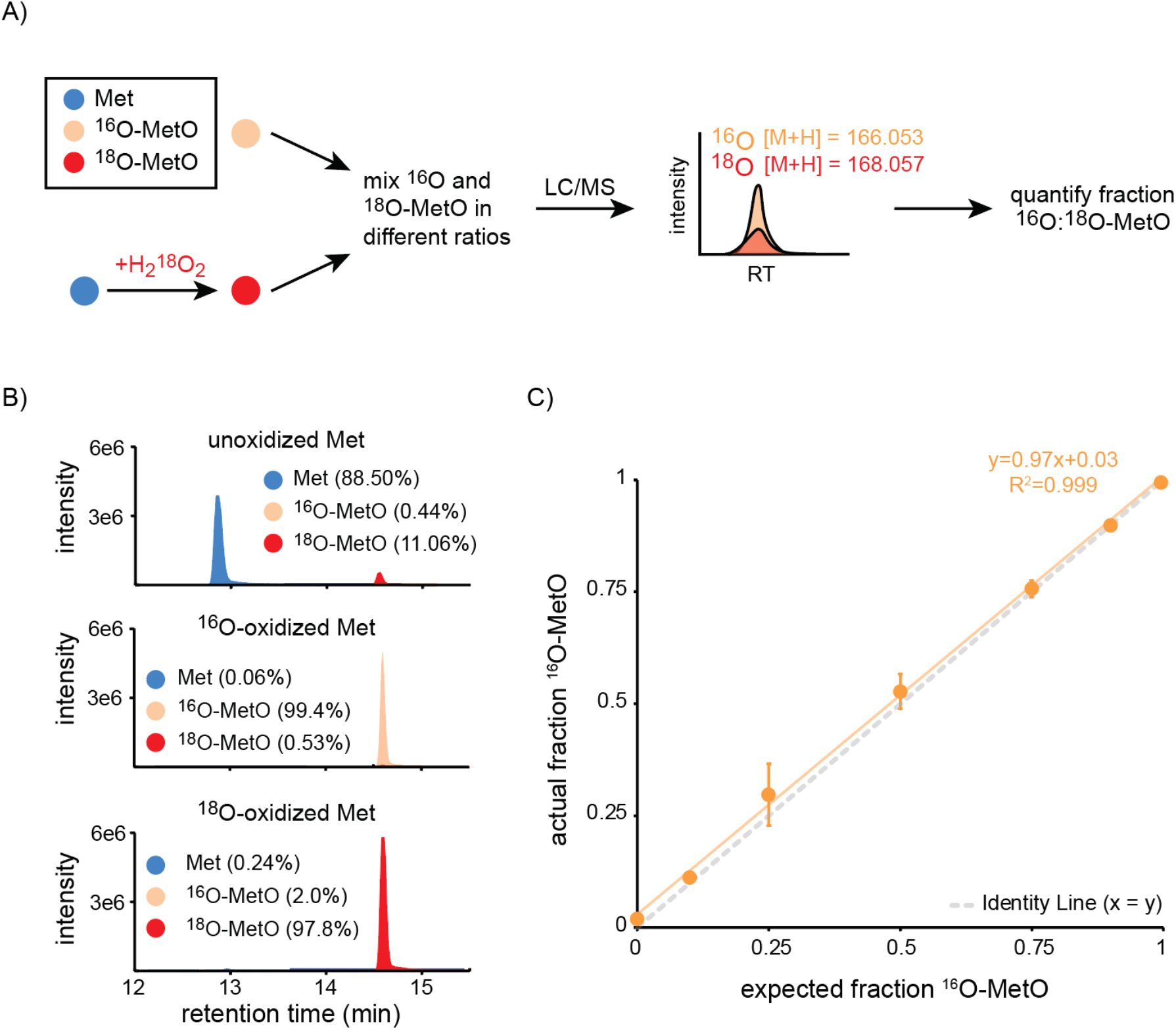
(A) Schematic of the workflow for assessing the accuracy of fMObB. (B) Relative levels of Met, ^16^O-MetO, and ^18^O-MetO in commercially obtained unoxidized Met and ^16^O-MetO, and in-house generated ^18^O-MetO samples. (C) Scatter plot indicating the correlation between measured and expected ratios of ^18^O-MetO to ^16^O-MetO in samples containing defined mixtures of the two.

### Free MetO levels in E. coli

We next applied fMObB to measure MetO levels in *E. coli* under varying conditions. Two different strains of *E. coli* were used in the analysis: a wildtype (WT) reference strain (MB6535) and a previously established mutant strain in which all 5 known bacterial MSRs had been deleted (MB6565, here referred to as -MSR).^29^ Using a data independent acquisition (DIA) proteomic workflow, we first verified that -MSR cells had depleted levels of MSR expression **(Fig. 2A**). Next, WT and -MSR strains were cultured under normal growth conditions or exposed to oxidative stress by the addition of non-lethal concentrations (100 μM) of hypochlorous acid (HOCl) for one hour. Extracts were obtained from the cells, and fractional oxidation of free Mets was measured by fMObB in triplicate measurements **(Fig. 2 B-C)**.

**Figure 2.**
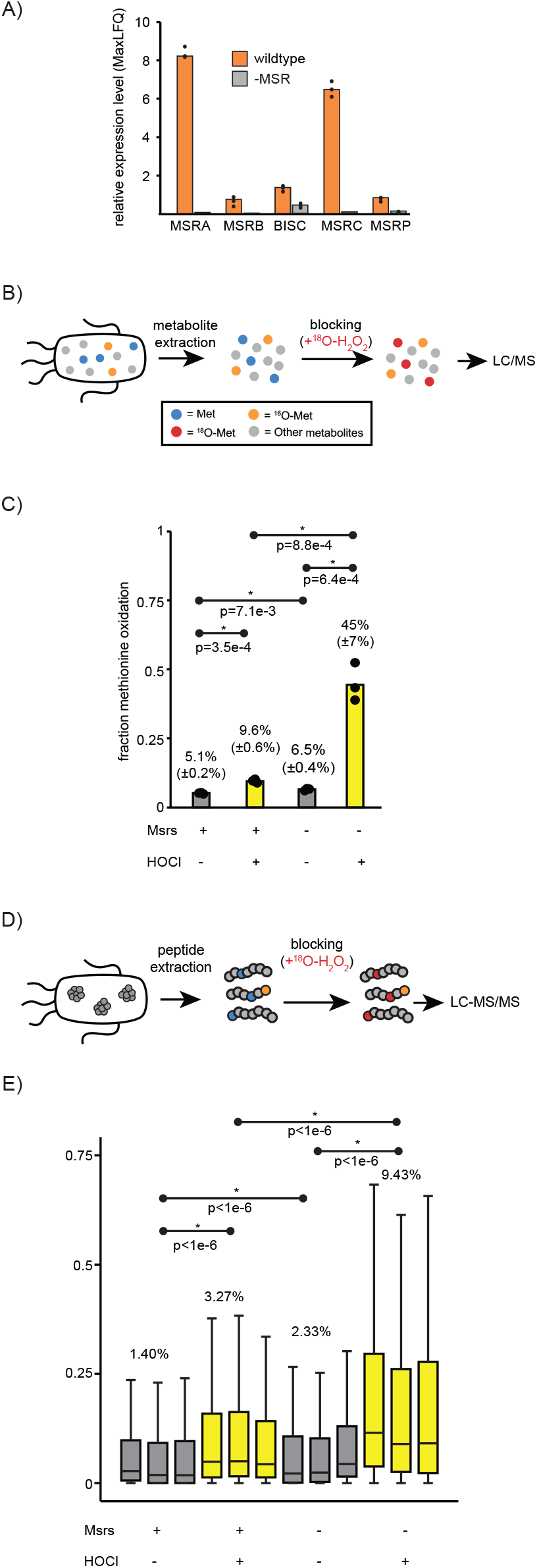
(A) Relative expression levels of MSRA, MSRB, BISC, MSRC and MSRP in unstressed wildtype an -MSR *E. coli* based on MaxLFQ measurements. (B) Schematic of the fMObB methodology for measurement of free methionine oxidation levels. (C) Fractional oxidation levels of free methionines in wildtype and -Msr *E. coli* in the absence and presence of hypochlorous acid (+HOCl). Medians of three replicate measurements are shown. p-values were calculated using two-tailed, equal variance t-test. (D) Schematic of the MObB methodology for measurement of protein-bound methionine oxidation levels. (E) Distributions of fractional oxidation levels of protein-bound methionines in wildtype and -Msr *E. coli* in the absence and presence of hypochlorous acid (+HOCl). The box plots indicate the range, medians, and interquartile range of three replicate experiments. p-values were calculated by applying the Mann– Whitney U test to the average of replicate experiments across the four conditions. The indicated percentages are the average percentage of methionines that are oxidized in each condition.

Under basal growth conditions, WT cells exhibited a free methionine oxidation level of 5.1% (±0.2%). In contrast, cells lacking MSRs displayed a significantly higher basal MetO level of 6.5% (±0.4%) (two-tailed, equal-variance t-test, *p* = 0.007). Exposure to HOCl resulted in a marked increase in methionine oxidation in both strains, with free MetO levels rising to 9.6%(0.6%) in WT cells and 45% (±7%) in the MSR-deficient strain. Together, these data establish a basal free methionine oxidation level of approximately 5% in *E. coli* and demonstrate that oxidative stress dramatically elevates free methionine oxidation. Moreover, the pronounced increase observed in the absence of MSRs indicates that, collectively, MSRs play a substantial role in maintaining reduced free methionine pools, particularly under conditions of oxidative stress.

### Protein-bound MetO levels in E. coli

We next evaluated how the oxidation levels of free Mets in *E. coli* compare to protein-bound Mets **(Fig. 2 D-E)**. Protein extracts from the experiments described above were analyzed using the previously described MobB protocol^6^ to globally measure the oxidation of Mets throughout the proteome. Between three replicate experiments, we were able to measure the oxidation levels of ~8,000 methionine residues mapped to ~2,400 proteins. Distributions of measured oxidation levels are plotted in **Fig. 2E**. Similar to the fMObB analyses above, we observed that oxidative stress and deletion of MSRs significantly increase the fractional oxidation of protein-bound Met residues. However, these increases were significantly less pronounced than free Mets. Under oxidative stress conditions in the -MSR strain, the average oxidation level of Met residues in the proteome was 9.4%.

Together, these results indicate that oxidative stress and loss of MSR activity lead to substantially higher oxidation levels of free Mets than of protein-bound Mets. These observations raise several important questions that warrant further investigation. First, it remains unclear whether the elevated oxidation of free Mets relative to protein-bound Mets reflects possible differences in intrinsic susceptibility to oxidation, a stronger dependence on MSR-mediated reduction, or formation from degradation of oxidized proteins. Second, the relative contributions of individual MSRs to the reduction of free versus protein-bound methionine sulfoxides under oxidative stress conditions remain unresolved. It is known that some MSRs exhibit higher *in vitro* catalytic efficiencies toward protein-bound methionine sulfoxides, whereas others preferentially reduce free methionine sulfoxides. For example, MSRC acts exclusively on free Met-R-O and shows among the highest *in vitro* activities recorded for any MSR against this substrate.^30^ However, how these specificities translate to net cellular activity is not well understood. As highlighted by our proteomic analyses (**Fig. 2A**), different MSRs are expressed at markedly different levels in *E. coli*, suggesting that their relative cellular abundances may strongly influence their overall contributions to methionine redox homeostasis. Future studies applying fMObB and MobB to strains lacking individual MSRs, or defined combinations thereof, will help clarify the division of labor within this enzyme family. More broadly, the fMObB approach established here provides a generalizable framework for probing the redox state of free methionine pools across diverse biological systems.

## Methods

### Preparation of ^16^O and ^18^O-MetO standards

^16^O-MetO stocks were prepared by dissolving L-MetO (Sigma) in water to a final concentration of 100 mM. To generate ^18^O-MetO, L-methionine (Fisher) was dissolved in water to a concentration of 200 mM and oxidized with 0.25 M H_2_^18^O_2_ (Sigma) for 30 minutes at 37°C, then frozen and lyophilized. The resulting ^18^O-MetO was brought up to a final concentration of 100 mM in water. Before mixing these two MetO species to generate the standard curve, ^16^O- and ^18^O-MetO stocks were diluted to 10 mM working concentrations. These working stocks were then mixed in different ratios (0:100, 10:90, 25:75, 50:50, 75:25, 90:10, 100:0) of ^16^O:^18^O-MetO in a final 20 μL volume.

### Bacteria culture and treatment

An E. coli strain derivative from MC4100 auxotroph for methionine (ΔmetB) lacking the genes for the five known prokaryotic MSRs – MsrA, MsrB, MsrC, BisC, and MsrP – named MB6565^29^, and its reference strain MC4100 ΔmetB (named JB590^31^) were streaked onto LB agar plates and incubated overnight at 37°C. Single colonies from these plates were used to inoculate a 5 mL culture in LB media, which were grown overnight at 30°C, shaking at 150 rpm due to the sensitivity of MB6565 to oxidative stress. The OD_600_ was taken of these overnight cultures before pelleting at 4000 xg for 5 minutes. Cells were washed with 1 mL 1x PBS (Corning) and spun down again at 4000 xg for 5 minutes. Cells were resuspended in 1x PBS to get an inoculum with an OD_600_ of 1. This resuspension was diluted 10x in 5 mL of minimal M9 media with 200 μM Met, inoculating two sets of three replicates for each strain. After 23 hours of growth, either 50 μL of 10 mM sodium hypochlorite (Sigma) or 50 μL 1X PBS was added and briefly vortexed prior to the final hour of incubation. The OD_600_ of the cultures was measured after this treatment period, and cultures were pelleted by centrifugation at 4000 xg for 5 minutes. Cells were washed twice with 1x PBS, pelleted again, and resuspended in PBS to obtain an OD_600_ of 1. For metabolomic analysis, 1 mL of the cell suspension was transferred to a new microcentrifuge tube, pelleted, and frozen in liquid nitrogen before storing at -80°C. The rest of the resuspension was pelleted in a separate microcentrifuge tube and frozen at -80°C before proteomic analysis.

### Metabolite extraction and blocking

Pellets for metabolomics were resuspended in 1 mL of 80:20 methanol to water metabolite extraction solution that was chilled on dry ice. Cells were kept on ice and lysed by sonication at 50% power, 15 seconds on and 30 seconds off for 3 cycles. The sample was transferred to dry ice for 15 minutes, then placed on regular ice for 30 minutes with a 5-second vortex every 10 minutes. Metabolite extract was separated from other cellular materials through a 10-minute centrifugation at 19,000 xg at 4°C. 800 μL of the supernatant was transferred to a new microcentrifuge tube and lyophilized. Metabolite extracts were resuspended in 25 μL of 0.1% trifluoroacetic acid (TFA) water. 25 μL of 1% H_2_^18^O_2_ solution, prepared by diluting 3% H_2_^18^O_2_ in 0.1% TFA water, was added to reach a final concentration of 0.5% H_2_^18^O_2_ (v/v) in each sample. Samples were oxidized by incubating at 37°C for 30 minutes with H_2_^18^O_2_ , then frozen at -80°C and lyophilized. Samples were reconstituted in 50% acetonitrile (A955, Fisher Scientific) and transferred to glass vials for LC/MS analysis.

### Mass spectrometry of metabolite extracts

For LC-MS analysis, metabolite extracts were analyzed by high-resolution mass spectrometry with an Orbitrap Exploris 240 (Thermo) coupled to a Vanquish Flex liquid chromatography system (Thermo). Samples were injected on a Waters XBridge XP BEH Amide column (150 mm length × 2.1 mm id, 2.5 μm particle size) maintained at 25°C, with a Waters XBridge XP VanGuard BEH Amide (5 mm × 2.1 mm id, 2.5 μm particle size) guard column. Mobile phase A was 100% LC-MS grade H2O with 10 mM ammonium formate and 0.125% formic acid. Mobile phase B was 90% acetonitrile with 10 mM ammonium formate and 0.125% formic acid. The gradient was 0 minutes, 100% B; 2 minutes, 100% B; 3 minutes, 90% B; 5 minutes, 90% B; 6 minutes, 85% B; 7 minutes, 85% B; 8 minutes, 75% B; 9 minutes, 75% B; 10 minutes, 55% B; 12 minutes, 55% B; 13 minutes, 35%, 20 minutes, 35% B; 20.1 minutes, 35% B; 20.6 minutes, 100% B; 22.2 minutes, 100% B all at a flow rate of 150 μl min^−1^, followed by 22.7 minutes, 100% B; 27.9 minutes, 100% B at a flow rate of 300 μl min_−1_, and finally 28 minutes, 100% B at flow rate of 150 μl min^−1^, for a total length of 28 minutes. The H-ESI source was operated in positive mode at spray voltage 3500 with the following parameters: sheath gas 35 au, aux gas 7 au, sweep gas 0 au, ion transfer tube temperature 320°C, vaporizer temperature 275° C, full scan MS1 mass resolution of 120,000 FWHM, RF lens at 70%, and standard automatic gain control (AGC).

### Proteomic sample preparation and blocking

Pellets collected for proteomic analysis were lysed with 50 μL of lysis buffer (5% SDS in 50 mM triethylammonium bicarbonate (TEAB) by sonication. Sonication was performed at 4°C in 30-second intervals at 50% power three times with 30-second breaks on ice in between. Samples were spun at 19,000 xg for 15 minutes at 4°C, and the supernatant was saved. Protein concentrations were determined by bicinchoninic acid (BCA) assay (Thermo). 25 μg of each sample was transferred to a new 0.5 mL tube, and 5% SDS in 50 mM TEAB was added to reach a total 25 μL volume. Samples were first incubated at 55°C for 60 min with 2.2 μL of 25 mM dithiothreitol (DTT). Samples were then incubated with 2.4 μL of 125 mM iodoacetamide (IAA) in the dark at room temperature for 20 minutes. Subsequently, 3 μL of 12% phosphoric acid was added to each sample along with 195.6 μL of 90% methanol in 100 mM TEAB. Samples were then added 165 μl at a time to S-Trap™ columns, spinning down the sample at 4,000 xg for 1 min in between each addition, and between each subsequent step. Columns were washed twice with 150 μL 90% methanol in 100 mM TEAB twice. 20 μL of 50 ng/uL trypsin was added to each column and allowed to soak for 5 minutes at room temperature before another 20 μL of 50 mM TEAB was added to the column. Samples were incubated at 37°C overnight in a water bath.

After incubation, peptides were eluted first with 40 μL of 0.1% trifluoroacetic acid (TFA) in water and then with 50/50 acetonitrile (ACN): 0.1% TFA water. Samples were frozen at -80°C and then lyophilized. Dried peptides were resuspended in 25 μL of 0.1% trifluoroacetic acid (TFA) water. 25 μL of 1% H_2_^18^O_2_ solution, prepared by diluting 3% H_2_^18^O_2_ in 0.1% TFA water, was added for a final concentration of 0.5% H_2_^18^O_2_ (v/v) in each sample. Samples were oxidized by incubating at 37°C for 30 minutes, then frozen at -80°C and lyophilized.

### Fractionation of proteomic samples

For MobB samples, dried down peptides were resuspended in 50 μL of 100 mM ammonium hydroxide (AH). Elution buffers for fractionation were made following the protocol below (Table 1). C18 stage tips were first activated by washing with acetonitrile (ACN) twice and then washing twice with 10 mM AH. Samples were loaded onto the stage tip and then washed once more with 10 mM AH. Fractions 1-8 were eluted with the elution buffer of the corresponding elution number. Every i and i+8 fractions (i.e. fraction 1 and fraction 9) were combined. Each wash and elution step was performed with 50 uL volumes by spinning at 2000 xg for 2 minutes. Fractions were frozen at -80°C and lyophilized. Fractionated samples were brought up in 0.1% TFA in water to 0.25 μg/μl and added to a 96-well sample plate.

**Table 1.** Details of elution buffers used for fractionation.

| **Elution Buffer** | **ACN (%)** | **ACN (uL)** | **10 mM AH (uL)** |
| --- | --- | --- | --- |
| E1 | 2 | 20 | 980 |
| E2 | 3.5 | 35 | 965 |
| E3 | 5 | 50 | 950 |
| E4 | 6.5 | 65 | 935 |
| E5 | 8 | 80 | 920 |
| E6 | 9.5 | 95 | 905 |
| E7 | 11 | 110 | 890 |
| E8 | 12.5 | 124 | 875 |
| E8 | 14 | 140 | 860 |
| E10 | 15.5 | 155 | 845 |
| E11 | 17 | 170 | 830 |
| E12 | 18.5 | 185 | 815 |
| E13 | 20 | 200 | 800 |
| E14 | 21.5 | 215 | 785 |
| E15 | 27 | 270 | 730 |
| E16 | 50 | 500 | 500 |

### Mass spectrometry of proteomic samples

Peptides were injected onto a 75 μm x 2 cm trap column (Thermo Fisher) prior to re-focusing on an Aurora Elite 75 μm x 15 cm C18 column (IonOpticks) using a Vanquish Neo UHPLC (Thermo Fisher), connected to an Orbitrap Astral mass spectrometer (Thermo Fisher). Solvent A was 0.1% formic acid in water, while solvent B was 0.1% formic acid in 80% acetonitrile. Ions were introduced to the mass spectrometer using an Easy-Spray source operating at 2 kV.

For samples analyzed using data-independent acquisition (DIA), the gradient began at 1% B and ramped to 5% B in 0.1 minutes, increased to 30% B in 12.1 minutes, increased to 40% in 0.7 minutes, and finally increased to 99% B in 0.1 minutes and was held for 2 minutes to wash the column for a total runtime of 15 minutes. After each run was completed, the column was re-equilibrated with 1% B prior to the next injection. The mass spectrometer was operated in DIA mode, with MS1 scans acquired in the Orbitrap at a resolution of 240,000, with a maximum injection time of 5 ms over a range of 380-980 *m/z*. DIA MS2 scans were acquired in the Astral mass analyzer with a 3 ms maximum injection time using a variable windowing scheme, using 2 Da windows from 380-680 *m/z*, 4 Da windows from 680-800 *m/z*, and 8 Da windows from 800-980 *m/z*. The HCD collision energy was set to 25%, and the normalized AGC was set to 500%. Fragment ions were collected over a scan range of 150-2000 *m/z*. The cycle time was 0.6 seconds.

For samples analyzed using data-dependent acquisition (DDA), the gradient began at 1% B and ramped to 5% B in 0.1 minutes, increased to 30% B in 21.3 minutes, increased to 40% in 1.5 minutes, and finally increased to 99% B in 0.1 minutes and was held for 2 minutes to wash the column for a total runtime of 25 minutes. After each run was completed, the column was re-equilibrated with 1% B prior to the next injection. The mass spectrometer was operated in DDA mode, with MS1 scans acquired in the Orbitrap and MS2 scans acquired in the Astral. The cycle time was set to 0.5 seconds. Monoisotopic Precursor Selection (MIPS) was set to “Peptide”. The full scan was done over a range of 375-1400 *m/z* with a resolution of 120,000 at *m/z* of 200, a normalized AGC target of 300%, and a maximum injection time of 10 ms. Peptides with a charge state between 2-5 were picked for fragmentation. Precursor ions were fragmented by high-energy collisional dissociation (HCD) using a collision energy of 28% with an isolation width of 1.1 *m/z*. The fragment scans were done over a range of 125-2000 *m/z* with a normalized AGC target of 300% and a maximum injection time of 10 ms. Dynamic exclusion was set to exclude after 1 time with a 6 second duration and a high and low mass tolerance of 7 ppm. The minimum filter intensity threshold was set to 5,000.

### Data analysis

For fMObB samples, LC-MS data were analyzed by Maven software for peak area determination, and metabolites were identified by matching to within-run standards. The spectra for each of the three metabolites of interest (Met, ^16^O-MetO, ^18^O-MetO) were specifically isolated for peak area determination by searching for their *m/z* in the protonated state (150.058, 166.053, and 168.057, respectively). The fraction oxidation in fMObB samples was determined by dividing the calculated intensity of ^16^O-MetO by the sum of the ^16^O-MetO and ^18^O-MetO intensity and averaging across three replicates.

For expression analysis of DIA data, the raw data were processed with DIA-NN version 1.9.2 (https://github.com/vdemichev/DIA-NN)^32^. For all experiments, data analysis was carried out using library-free analysis mode in DIA-NN. To annotate the library, the *E. coli* strain K-12 UniProt ‘one protein sequence per gene’ database (UP000000625, downloaded 02/10/2025) was used, with ‘deep learning-based spectra and RT prediction’ enabled. For precursor ion generation, the maximum number of missed cleavages was set to 1, maximum number of variable modifications to 1 for Ox(M), peptide length range to 7-30, precursor charge range to 2-3, precursor *m/z* range to 380-980, and fragment *m/z* range to 150-2000. The quantification was set to ‘Robust LC (high precision)’ mode with normalization set to RT-dependent, MBR enabled, protein inferences set to ‘Genes’, and ‘Heuristic protein inference’ turned off. MS1 and MS2 mass tolerances, along with the scan window size were automatically set by the software. Precursors were subsequently filtered at library precursor q-value (1%), library protein group q-value (1%), and posterior error probability (50%). Protein quantification was carried out using the MaxLFQ intensities obtained from the DIA-NN output files which were averaged across three replicates of the WT strain.

Measurement of protein-bound methionine oxidation levels by MObB were conducted using a DDA workflow as described above. Raw data were searched using MSFragger v 4.1 in FragPipe v. 22.0^33^. The search in FragPipe used a Uniprot *E. coli* database (UP000000625_8333, downloaded 05/05/2025) supplemented with decoys and contaminants by FragPipe. Methionine oxidation with ^16^O and ^18^O were set as variable modifications along with N-terminal acetylation, and carbamidomethylated cysteine was set as a fixed modification. A maximum of 3 modifications was allowed per peptide. The precursor mass tolerance was set at 20 ppm. Mass calibration and parameter optimization were enabled along with validation tools. Trypsin was set as the protease with up to 2 missed cleavages allowed. MS1 files were generated from raw files using MSConvert^34^. MS1 files and Fragpipe-generated.psm files were used to measure fractional oxidation levels as previously described^6,25^. All raw data and search results are available in the ProteomeXchange Consortium via the PRIDE partner repository (accession number PXD076670).^35,36^

## Author Information

### Author Contributions

The study concept was conceived by M.A., P.B. and S.G. The experiments were carried out by M.A. Mass spectrometry was performed by K.W., K.S. and J.H. Data analysis was conducted by M.A., and S.G. Resources were contributed by B.M. The initial draft of the manuscript was written by M.A. and S.G. All authors have given approval to the final version of the manuscript.

## Abbreviations

Met: methionine
MetO: methionine sulfoxide
ROS: reactive oxygen species
MSR: methionine sulfoxide reductase
MSRA: methionine sulfoxide reductase A
MSRB: methionine sulfoxide reductase B
MobB: Methionine oxidation by Blocking
fMObB: free Methionine Oxidation by Blocking
DIA: data independent acquisition
HOCl: hypochlorous acid
BEH: bridged ethyl hybrid
H-ESI: heated electrospray ionization
AGC: automatic gain control
TEAB: triethylammonium bicarbonate
FWHM: full-width at half maximum
RF: radio frequency
BCA: bicinchoninic acid
DTT: dithiothreitol
IAA: iodoacetamide
TFA: trifluoroacetic acid
ACN: acetonitrile
AH: ammonium hydroxide
HCD: higher-energy collisional dissociation
DDA: data dependent acquisition
MIPS: Monoisotopic Precursor Selection
MBR: match-between-runs
MaxLFQ: maximal label-free quantification

## Acknowledgements

We would like to thank the Metabolomics and Mass Spectrometry Resource Laboratories at the University of Rochester Medical Center (URMC) for collecting the mass spectrometry data. We would also like to thank the Beckman Foundation (Beckman Scholars Program) for financially supporting this work.

## References

(1) Manta, B.; Gladyshev, V. N. Regulated Methionine Oxidation by Monooxygenases. Free Radic. Biol. Med. 2017, 109, 141–155. 10.1016/j.freeradbiomed.2017.02.010.

(2) Ezraty, B.; Aussel, L.; Barras, F. Methionine Sulfoxide Reductases in Prokaryotes. Biochim. Biophys. Acta BBA-Proteins Proteomics 2005, 1703 (2), 221–229. 10.1016/j.bbapap.2004.08.017.

(3) Jiang, B.; Moskovitz, J. The Functions of the Mammalian Methionine Sulfoxide Reductase System and Related Diseases. Antioxidants 2018, 7 (9), 122. 10.3390/antiox7090122.

(4) Stadtman, E. R.; Van Remmen, H.; Richardson, A.; Wehr, N. B.; Levine, R. L. Methionine Oxidation and Aging. Biochim. Biophys. Acta 2005, 1703 (2), 135–140. 10.1016/j.bbapap.2004.08.010.

(5) Butterfield, D. A.; Boyd-Kimball, D. The Critical Role of Methionine 35 in Alzheimer’s Amyloid β-Peptide (1–42)-Induced Oxidative Stress and Neurotoxicity. Biochim. Biophys. Acta BBA-Proteins Proteomics 2005, 1703 (2), 149–156. 10.1016/j.bbapap.2004.10.014.

(6) Bettinger, J. Q.; Simon, M.; Korotkov, A.; Welle, K. A.; Hryhorenko, J. R.; Seluanov, A.; Gorbunova, V.; Ghaemmaghami, S. Accurate Proteomewide Measurement of Methionine Oxidation in Aging Mouse Brains. J. Proteome Res. 2022, 21 (6), 1495–1509. 10.1021/acs.jproteome.2c00127.

(7) Luo, S.; Levine, R. L. Methionine in Proteins Defends against Oxidative Stress. FASEB J. 2009, 23 (2), 464–472. 10.1096/fj.08-118414.

(8) Campbell, K.; Vowinckel, J.; Keller, M. A.; Ralser, M. Methionine Metabolism Alters Oxidative Stress Resistance via the Pentose Phosphate Pathway. Antioxid. Redox Signal. 2016, 24 (10), 543–547. 10.1089/ars.2015.6516.

(9) Drazic, A.; Miura, H.; Peschek, J.; Le, Y.; Bach, N. C.; Kriehuber, T.; Winter, J. Methionine Oxidation Activates a Transcription Factor in Response to Oxidative Stress. Proc. Natl. Acad. Sci. 2013, 110 (23), 9493–9498. 10.1073/pnas.1300578110.

(10) Anbanandam, A.; Bieber Urbauer, R. J.; Bartlett, R. K.; Smallwood, H. S.; Squier, T. C.; Urbauer, J. L. Mediating Molecular Recognition by Methionine Oxidation: Conformational Switching by Oxidation of Methionine in the Carboxyl-Terminal Domain of Calmodulin. Biochemistry 2005, 44 (27), 9486–9496. 10.1021/bi0504963.

(11) Walker, E. J.; Bettinger, J. Q.; Welle, K. A.; Hryhorenko, J. R.; Ghaemmaghami, S. Global Analysis of Methionine Oxidation Provides a Census of Folding Stabilities for the Human Proteome. Proc. Natl. Acad. Sci. 2019, 116 (13), 6081–6090. 10.1073/pnas.1819851116.

(12) Li, J.; Ge, P.; He, Q.; Liu, C.; Zeng, C.; Tao, C.; Zhai, Y.; Wang, J.; Zhang, Q.; Wang, R.; Zhang, Y.; Zhang, D.; Zhao, J. Association between Methionine Sulfoxide and Risk of Moyamoya Disease. Front. Neurosci. 2023, 17. 10.3389/fnins.2023.1158111.

(13) Cho, Y.; Park, Y.; Sim, B.; Kim, J.; Lee, H.; Cho, S.-N.; Kang, Y. A.; Lee, S.-G. Identification of Serum Biomarkers for Active Pulmonary Tuberculosis Using a Targeted Metabolomics Approach. Sci. Rep. 2020, 10, 3825. 10.1038/s41598-020-60669-0.

(14) Moskovitz, J.; Smith, A. Methionine Sulfoxide and the Methionine Sulfoxide Reductase System as Modulators of Signal Transduction Pathways: A Review. Amino Acids 2021, 53 (7), 1011–1020. 10.1007/s00726-021-03020-9.

(15) Free methionine-(R)-sulfoxide reductase from Escherichia coli reveals a new GAF domain function | PNAS. https://www.pnas.org/doi/10.1073/pnas.0703774104 (accessed 2026-02-13).

(16) Lin, Z.; Johnson, L. C.; Weissbach, H.; Brot, N.; Lively, M. O.; Lowther, W. T. Free Methionine-(R)-Sulfoxide Reductase from Escherichia Coli Reveals a New GAF Domain Function. Proc. Natl. Acad. Sci. 2007, 104 (23), 9597–9602. 10.1073/pnas.0703774104.

(17) Andrieu, C.; Vergnes, A.; Loiseau, L.; Aussel, L.; Ezraty, B. Characterisation of the Periplasmic Methionine Sulfoxide Reductase (MsrP) from Salmonella Typhimurium. Free Radic. Biol. Med. 2020, 160, 506–512. 10.1016/j.freeradbiomed.2020.06.031.

(18) Kim, J.-S.; Liu, L.; Kant, S.; Orlicky, D. J.; Uppalapati, S.; Margolis, A.; Davenport, B. J.; Morrison, T. E.; Matsuda, J.; McClelland, M.; Jones-Carson, J.; Vazquez-Torres, A. Anaerobic Respiration of Host-Derived Methionine Sulfoxide Protects Intracellular Salmonella from the Phagocyte NADPH Oxidase. Cell Host Microbe 2024, 32 (3), 411–424.e10. 10.1016/j.chom.2024.01.004.

(19) Javitt, G.; Cao, Z.; Resnick, E.; Gabizon, R.; Bulleid, N. J.; Fass, D. Structure and Electron-Transfer Pathway of the Human Methionine Sulfoxide Reductase MsrB3. Antioxid. Redox Signal. 2020, 33 (10), 665–678. 10.1089/ars.2020.8037.

(20) Mashima, R.; Nakanishi-Ueda, T.; Yamamoto, Y. Simultaneous Determination of Methionine Sulfoxide and Methionine in Blood Plasma Using Gas Chromatography-Mass Spectrometry. Anal. Biochem. 2003, 313 (1), 28–33. 10.1016/S0003-2697(02)00537-7.

(21) Potgieter, H. C.; Ubbink, J. B.; Bissbort, S.; Bester, M. J.; Spies, J. H.; Vermaak, W.J. H. Spontaneous Oxidation of Methionine: Effect on the Quantification of Plasma Methionine Levels. Anal. Biochem. 1997, 248 (1), 86–93. 10.1006/abio.1997.2075.

(22) Li, Z.; Majeski, J. B.; Choe, R.; Schmitthenner, H. F.; Baran, T. M. Localization of a Molecularly Targeted Probe for Imaging and Photodynamic Therapy of Breast Cancer in a Murine Model. In Optical Methods for Tumor Treatment and Detection: Mechanisms and Techniques in Photodynamic Therapy XXXI; SPIE, 2023; Vol. PC12359, p PC123590D. 10.1117/12.2648451.

(23) Zang, L.; Carlage, T.; Murphy, D.; Frenkel, R.; Bryngelson, P.; Madsen, M.; Lyubarskaya, Y. Residual Metals Cause Variability in Methionine Oxidation Measurements in Protein Pharmaceuticals Using LC-UV/MS Peptide Mapping. J. Chromatogr. B 2012, 895–896, 71–76. 10.1016/j.jchromb.2012.03.016.

(24) Chen, M.; Cook, K. D. Oxidation Artifacts in the Electrospray Mass Spectrometry of Aβ Peptide. Anal. Chem. 2007, 79 (5), 2031–2036. 10.1021/ac061743r.

(25) Bettinger, J. Q.; Welle, K. A.; Hryhorenko, J. R.; Ghaemmaghami, S. Quantitative Analysis of in Vivo Methionine Oxidation of the Human Proteome. J. Proteome Res. 2020, 19 (2), 624–633. 10.1021/acs.jproteome.9b00505.

(26) Rauh, M. Steroid Measurement with LC–MS/MS. Application Examples in Pediatrics. J. Steroid Biochem. Mol. Biol. 2010, 121 (3), 520–527. 10.1016/j.jsbmb.2009.12.007.

(27) Kondyli, A.; Schrader, W. Evaluation of the Combination of Different Atmospheric Pressure Ionization Sources for the Analysis of Extremely Complex Mixtures. Rapid Commun. Mass Spectrom. 2020, 34 (8), e8676. 10.1002/rcm.8676.

(28) Liu, H.; Ponniah, G.; Neill, A.; Patel, R.; Andrien, B. Accurate Determination of Protein Methionine Oxidation by Stable Isotope Labeling and LC-MS Analysis. Anal. Chem. 2013, 85 (24), 11705–11709. 10.1021/ac403072w.

(29) Anton, B. P.; Morgan, R. D.; Ezraty, B.; Manta, B.; Barras, F.; Berkmen, M. Complete Genome Sequence of Escherichia Coli BE104, an MC4100 Derivative Lacking the Methionine Reductive Pathway. Microbiol. Resour. Announc. 2019, 8 (28), e00721–19. 10.1128/MRA.00721-19.

(30) Kryukov, G. V.; Wilson, F. H.; Ruth, J. R.; Paulk, J.; Tsherniak, A.; Marlow, S. E.; Vazquez, F.; Weir, B. A.; Fitzgerald, M. E.; Tanaka, M.; Bielski, C. M.; Scott, J. M.; Dennis, C.; Cowley, G. S.; Boehm, J. S.; Root, D. E.; Golub, T. R.; Clish, C. B.; Bradner, J. E.; Hahn, W. C.; Garraway, L. A. MTAP Deletion Confers Enhanced Dependency on the PRMT5 Arginine Methyltransferase in Cancer Cells. Science 2016, 351 (6278), 1214–1218. 10.1126/science.aad5214.

(31) Gennaris, A.; Ezraty, B.; Henry, C.; Agrebi, R.; Vergnes, A.; Oheix, E.; Bos, J.; Leverrier, P.; Espinosa, L.; Szewczyk, J.; Vertommen, D.; Iranzo, O.; Collet, J.-F.; Barras, F. Repairing Oxidized Proteins in the Bacterial Envelope Using Respiratory Chain Electrons. Nature 2015, 528 (7582), 409–412. 10.1038/nature15764.

(32) Demichev, V.; Messner, C. B.; Vernardis, S. I.; Lilley, K. S.; Ralser, M. DIA-NN: Neural Networks and Interference Correction Enable Deep Proteome Coverage in High Throughput. Nat. Methods 2020, 17 (1), 41–44. 10.1038/s41592-019-0638-x.

(33) Kong, A. T.; Leprevost, F. V.; Avtonomov, D. M.; Mellacheruvu, D.; Nesvizhskii, A. I. MSFragger: Ultrafast and Comprehensive Peptide Identification in Shotgun Proteomics. Nat. Methods 2017, 14 (5), 513–520. 10.1038/nmeth.4256.

(34) Chambers, M. C.; Maclean, B.; Burke, R.; Amodei, D.; Ruderman, D. L.; Neumann, S.; Gatto, L.; Fischer, B.; Pratt, B.; Egertson, J.; Hoff, K.; Kessner, D.; Tasman, N.; Shulman, N.; Frewen, B.; Baker, T. A.; Brusniak, M.-Y.; Paulse, C.; Creasy, D.; Flashner, L.; Kani, K.; Moulding, C.; Seymour, S. L.; Nuwaysir, L. M.; Lefebvre, B.; Kuhlmann, F.; Roark, J.; Rainer, P.; Detlev, S.; Hemenway, T.; Huhmer, A.; Langridge, J.; Connolly, B.; Chadick, T.; Holly, K.; Eckels, J.; Deutsch, E. W.; Moritz, R. L.; Katz, J. E.; Agus, D. B.; MacCoss, M.; Tabb, D. L.; Mallick, P. A Cross-Platform Toolkit for Mass Spectrometry and Proteomics. Nat. Biotechnol. 2012, 30 (10), 918–920. 10.1038/nbt.2377.

(35) Perez-Riverol, Y.; Bandla, C.; Kundu, D. J.; Kamatchinathan, S.; Bai, J.; Hewapathirana, S.; John, N. S.; Prakash, A.; Walzer, M.; Wang, S.; Vizcaíno, J. A. The PRIDE Database at 20 Years: 2025 Update. Nucleic Acids Res. 2025, 53 (D1), D543–D553. 10.1093/nar/gkae1011.

(36) Deutsch, E. W.; Bandeira, N.; Perez-Riverol, Y.; Sharma, V.; Carver, J. J.; Mendoza, L.; Kundu, D. J.; Wang, S.; Bandla, C.; Kamatchinathan, S.; Hewapathirana, S.; Pullman, B. S.; Wertz, J.; Sun, Z.; Kawano, S.; Okuda, S.; Watanabe, Y.; MacLean, B.; MacCoss, M. J.; Zhu, Y.; Ishihama, Y.; Vizcaíno, J. A. The ProteomeXchange Consortium at 10 Years: 2023 Update. Nucleic Acids Res. 2023, 51 (D1), D1539–D1548. 10.1093/nar/gkac1040.

